# New insights into homoeologous copy number variations in the hexaploid wheat genome

**DOI:** 10.1101/2020.09.09.289447

**Authors:** Caroline Juery, Lorenzo Concia, Romain De Oliveira, Nathan Papon, Ricardo Ramírez-González, Moussa Benhamed, Cristobal Uauy, Frédéric Choulet, Etienne Paux

## Abstract

Bread wheat is an allohexaploid species originating from two successive and recent rounds of hybridization between three diploid species that were very similar in terms of chromosome number, genome size, TE content, gene content and synteny. As a result, it has long been considered that most of the genes were in three pairs of homoeologous copies. However, these so-called triads represent only one half of wheat genes, while the remaining half belong to homoeologous groups with various number of copies across subgenomes. In this study, we examined and compared the distribution, conservation, function, expression and epigenetic profiles of triads with homoeologous groups having undergone a deletion (dyads) or a duplication (tetrads) in one subgenome. We show that dyads and tetrads are mostly located in distal regions and have lower expression level and breadth than triads. Moreover, they are enriched in functions related to adaptation and more associated with the repressive H3K27me3 modification. Altogether, these results suggest that triads mainly correspond to housekeeping genes and are part of the core genome, while dyads and tetrads belong to the *Triticeae* dispensable genome. In addition, by comparing the different categories of dyads and tetrads, we hypothesize that, unlike most of the allopolyploid species, subgenome dominance and biased fractionation are absent in hexaploid wheat. Differences observed between the three subgenomes are more likely related to two successive and ongoing waves of post-polyploid diploidization, that had impacted A and B more significantly than D, as a result of the evolutionary history of hexaploid wheat.

**Core ideas:**

Only one half of hexaploid wheat genes are in triads, *i.e*. in a 1:1:1 ratio across subgenomes
Triads are likely part of the core genome; dyads and tetrads belong to the dispensable genome
Subgenome dominance and biased fractionation are absent in hexaploid wheat
Subgenome differences are related to two successive waves of post-polyploid diploidization

## Introduction

Polyploidy, or whole genome duplication (WGD), has been observed in many living organisms, both prokaryotic and eukaryotic, and is widely recognized as a key driver of species evolution, diversification, as well as domestication (Van de Peer et al., 2017; Wendel, 2015). Polyploid species can be classified into two different categories: autopolyploids, which arise from genome doubling within one species, and allopolyploids, which arise from genome doubling following hybridization between two distinct species (Glover et al., 2016). About one half of angiosperms are recent polyploids, including numerous important crop species such as oilseed rape (*Brassica napus*, 7500 years old), coffee (*Coffea arabica*, 10,000-50,000 years old) and wheat (*Triticum aestivum*, 10,000 years old). In addition, due to recurrent polyploidization events that occured through time, including the ζ (zeta) WGD event 300-350 million years ago (MYA), all flowering plants are ancient polyploids or paleopolyploids (Jiao et al., 2011). Well-characterized examples include sorghum (*Sorghum bicolor*, 95-115 MYA), maize (*Zea mays*, 26 MYA) and soybean (*Glycine max,13* MYA) (Qiao et al., 2019).

Each WGD event results in a doubling of the gene content. Fang and Morrell (2016) demonstrated that polyploidization gives fitness advantages through increasing the amount of raw genetic material on which natural and artificial selection can happen. However, despite the successive episodes of WGDs that occurred through time, the number of genes in plants is quite similar (Michael and Jackson, 2013) and far less than that expected by the doubling process (Adams and Wendel, 2005). This leads to the paradox that while being an important evolutionary process, polyploidy also seems to be an evolutionary ‘dead-end’ (Van de Peer et al., 2017) as polyploids systematically tend to return to a diploid state after a few million years (Wendel, 2015). Hence, duplicated genes can either be pseudogenized, silenced, and eventually lost or, alternatively, retained because having evolved a new function (neofunctionalization) or having diverged in expression (subfunctionalization) (Flagel and Wendel, 2009).

Previous studies on homoeologous gene loss and retention, as well as relative expression contribution in various polyploid species, revealed species-specific patterns, suggesting an effect of the age of the polyploidization and diploid progenitor divergence (Bottani et al., 2018). For example, in the paleopolyploid maize genome, 14% of coding sequences were lost during the diploidization process, with a 25% of differential loss between the two genomes and a biased fractionation (loss of functioning DNA sequence) in favor of one subgenome that exhibits an overall higher expression and higher impact on phenotypic variability (Jiao et al., 2017; Renny-Byfield et al., 2017). Similarly, in the ancient allotetraploid cotton genome, a biased fractionation was observed, with the A genome showing more gene loss, a faster evolution rate, and an overall lower expression level that the D genome (Zhang et al., 2015).

In contrast, while frequent homoeolog sequence exchanges have been reported, no significant bias toward either subgenome was observed in the recent allotetraploid oilseed rape (*Brassica napus*) (Chalhoub et al., 2014). Therefore, for closely related progenitor genomes, like in soybean, a dosage sensitive pattern of expression leads to stochastic differentiation of homoeologous pairs. For highly divergent progenitor genomes, like maize, the more favorable homoeologous genes set of a subgenome are selected, leading to an overall subgenome retention and a biased fractionation.

Bread wheat (*Triticum aestivum* L.) is an allohexaploid species (2n = 6X = AABBDD) originating from two successive rounds of hybridization (International Wheat Genome Sequencing Consortium, 2014; Marcussen et al., 2014). The first hybridization event occurred ~800,000 years ago between *Triticum urartu* (AA-genome) and an unknown *Aegilops* species (BB-genome). The second event took place ~10,000 years ago between *Triticum turgidum* (AABB-genome) and *Aegilops tauschii* (DD-genome). The resulting hexaploid AABBDD-genome was estimated to carry 107,891 high confidence (HC) protein-coding genes, although 161,537 low confidence (LC) genes and 303,818 pseudogenes and gene fragments were also annotated (International Wheat Genome Sequencing Consortium, 2018). Using a phylogenomics approach on a filtered set of 181,036 genes, 21,603 triads, defined as homoeologous genes that had a strict 1:1:1 correspondence (one copy per subgenome A, B and D), were identified. These account for only 36% of the gene set (64,809 genes) while the remaining 64% correspond have a more complex homoeologous relationships (1:1:N or 0:1:1 for example). Similar proportions of genes in different homoeology contexts were observed on each of the subgenome. Together with equal contribution of the three homoeologous genomes to the overall gene expression, this supported the hypothesis of the absence of biased fractionation and global subgenome dominance (International Wheat Genome Sequencing Consortium, 2018). However, a cell type- and stage-dependent local subgenome dominance was observed (Harper et al., 2016; Pfeifer et al., 2014). A recent study reported that the vast majority of triads displayed a balanced contribution of each copy to the overall expression of the homoeologous group (Ramirez-Gonzalez et al., 2018). For those showing either dominance or suppression of one homoeologous copy, differences were associated with epigenetic changes, especially in H3K9ac and H3K27me3 patterns. Such differences in gene expression likely represent the first steps toward neo- or subfunctionalization of wheat homoeologs.

Previous studies focused mainly on 1:1:1 triads leaving two third of the wheat genes apart. However, triads likely correspond to highly conserved and evolutionary constrained genes. In this regard, they may not be fully representative of the entire gene set and may not illustrate the complexity of the evolutionary trajectories that occurred within the hexaploid wheat genome. Here, we report on this unexplored part of the wheat genome by integrating not only triads but also dyads and tetrads, *i.e*. homoeologous groups that have undergone a single gene loss or duplication event, respectively. By combining genomic, transcriptomic and epigenetic data, we show that these two latter categories differ from triads not only by their chromosomal distribution but also by their transcriptional and epigenetic patterns as well as their conservation in wheat and other plant genomes, suggesting different evolutionary fates depending on the copy number of homoeologous genes.

## Material and Methods

### Definition and distribution of homoeologous group

Dyad, triad and tetrad gene information were retrieved from IWGSC (2018). Groups containing both high-confidence (HC) and low-confidence (LC) genes were filtered out to keep only those with HC genes. These data included gene position on the IWGSC RefSeq v1.0 reference sequence, homoeologous group category (dyad, triad or tetrad) and ID, as well as orthologous relationships with *Arabidopsis thaliana, Zea mays, Sorghum bicolor, Oryza sativa, Brachypodium distachyon* and *Hordeum vulgare* genes. The chromosome distribution of genes was performed by calculating the proportion of dyad, triad and tetrad genes over the total number of genes from this study within each of the five chromosomal regions defined by Pingault et al. (2015). Duplicated genes separated by less than 10 genes and less than 1 Mb on chromosomes were considered as tandem duplications. The other ones were considered as dispersed duplications.

### Characterization of ancestral duplications / deletions and presence-absence variations

To assess if deletions in dyads occurred within diploids or upon polyploidization, we aligned the two remaining copies onto diploid and tetraploid ancestor genomes using the GMAP package v2019.03.04 (Wu and Watanabe, 2005) with 85 % of sequence identity and 85% of sequence coverage as parameters. For AB-dyads, we mined the *Aegilops tauschii* genome (Luo et al., 2017) with A and B coding sequences. For AD-dyads, we mined the B-genome of *Triticum dicoccoides* (Avni et al., 2017) with A and D coding sequences. For BD dyads, we mined the *Triticum urartu* genome (Ling et al., 2018), as well as the A-genome of *Triticum dicoccoides* with B and D sequences. To take into account polymorphisms between individual sequences (divergence, presence / absence variations...), we corrected these numbers by dividing them by the number of genes that are still present in the hexaploid wheat genome and that were found in the ancestral genomes (*e.g*. D-copy from an AD dyad in the *Ae. tauschii* genome). To assess if duplications in tetrads occurred within diploids or upon polyploidization, we used GMAP to estimate the number of copies in the diploid and tetraploid ancestor genomes. Presence/absence variation (PAV) analysis was performed as described by De Oliveira et al. (2020). Briefly, sequencing reads from 16 wheat accessions (Montenegro et al., 2017) were mapped on the IWGSC RefSeq v1.0 using BWA-MEM v0.7.12 (Li and Durbin, 2010). Alignments were then filtered out using samtools view (samtools view -F 2308 -q11; Li et al., 2009) and PCR duplicates were removed using samtools rmdup. Depth of coverage was assessed using bedtools coverage (v2.26; Quinlan and Hall, 2010). Genes were considered as putative PAVs when their coding sequence was covered over less than 10% of its length in at least two accessions.

### Gene Ontology enrichment and functional analysis

Gene Ontology (GO) terms and functional annotation data were retrieved from IWGSC (2018). GO enrichment analysis was conducted using R package *topGO* (Alexan and Rahnenfuhrer, 2019).

### Gene expression and relative contribution analysis

Expression data from 15 samples representing five different organs (root, leaf, stem, spike and grain) at three developmental stages each in controlled non-stressed conditions were retrieved from Ramirez-Gonzalez et al. (2018). Genes with expression levels below 0.5 TPM were considered as non-expressed. Outlier identification was conducted in R (R Core Team, 2014) using the *boxplot* function of the *ggplot2* package (Villanueva and Chen, 2019).

For relative contribution analyses, we used the calculation method described by Ramirez-Gonzalez et al. (2018). Briefly, to standardize the relative expression of each homoeolog across a group, we normalized the absolute TPM for each gene within this group, as follows:

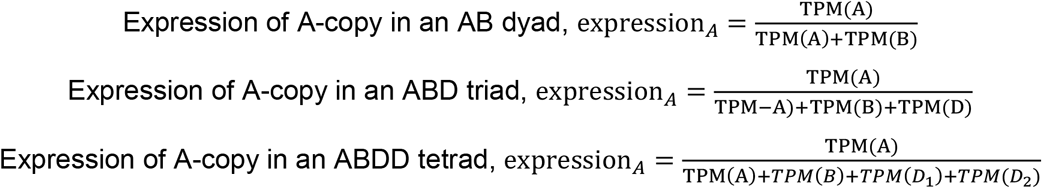

The normalized expression was calculated for the average across all expressed tissues as well as for each tissue individually. In order to assign theorical expression bias categories to each group within triad, tetrad and dyad, we constructed theorical matrix (Table S1). We calculated the Euclidean distance with the *rdist* function from R from the observed normalized expression of each group to each of the ideal categories. We assigned the homoeolog expression bias category for each group by selecting the shortest distance between theorical and observed relative contribution values. For binary organ expression, genes expressed in an organ (≥ 0.5 TPM) were given a value of 1 and those not expressed (<0.5 TPM), 0. This resulted in 32 binary expression profiles (0-0-0-0-0, 0-0-0-0-1, 0-0-0-1-1...).

### Histone mark analysis

Wheat H3K9ac and H3K27me3 data from bread wheat cultivar Chinese Spring at three-leaf stage were retrieved from IWGSC (2018). Genes were assigned a histone mark category, either H3K9ac, H3K27me3, both or no mark. We calculated meta-gene profiles for each category by computing the read density of each histone mark over different categories using Deeptools (Ramirez et al., 2016) computeMatrix scale-regions and plotted it with plotProfile. Only reads mapping within gene bodies plus 1kb upstream of the transcription start site and 1kb downstream of the transcription end site were considered. To account for different gene size, we divided the read counts over each gene by its length. H3K9ac and H3K27me3 data from *Oryza sativa* and *Zea mays* were retrieved from the Plant Chromatin State Database (Liu et al., 2018).

## Results

### Defining homoeologous groups

To decipher the impact of gene loss and duplication in the wheat genome, we focused our study on high confidence (HC) genes from the IWGSC RefSeq v1.0. Out of the 107,891 HC genes, we found 55,170 homoeologs (51.1%) belonging to 18,390 triads (Tables 1 and S2, Data S1). It is worth noting that an additional 2,218 triads containing both HC and LC genes were found. However, since LC genes corresponded to partially supported gene models, we did not include these triads in our analysis. We also found 12,640 genes (11.7%) corresponding to 6,320 groups having undergone a single gene loss during the course of evolution since the divergence of the A, B, and D subgenomes. The corresponding groups will be hereafter referred to as AB, AD and BD dyads, *i.e*. groups of HC genes being in 1:1:0, 1:0:1 or 0:1:1: ratios across homoeolog genomes, respectively. Finally, we identified 3,008 genes (2.8%) belonging to 240 AABD, 315 ABBD and 197 ABDD tetrads, *i.e*. groups having undergone one single gene duplication thus being in 2:1:1, 1:2:1 or 1:1:2 ratios, respectively. Similar to triads, 2,085 and 658 additional dyads and tetrads containing both HC and LC genes were found but not selected for further analyses. While in the present study we will focus only on dyads, triads and tetrads (70,818 genes), it is worth noting that 37,073 HC genes (34.4%) depart from these ratios, corresponding to genes that have undergone more than one deletion or duplication or that were not clustered into a homoeolog group. The overall high proportion of genes that are not in a strict 1:1:1 ratio, hereafter referred to as genes affected by homoeologous copy number variations (HomoeoCNVs), represent the dynamic part of the wheat genome during the course of its evolution either in the ancestral diploid species or after polyploidization.

**Table 1.**
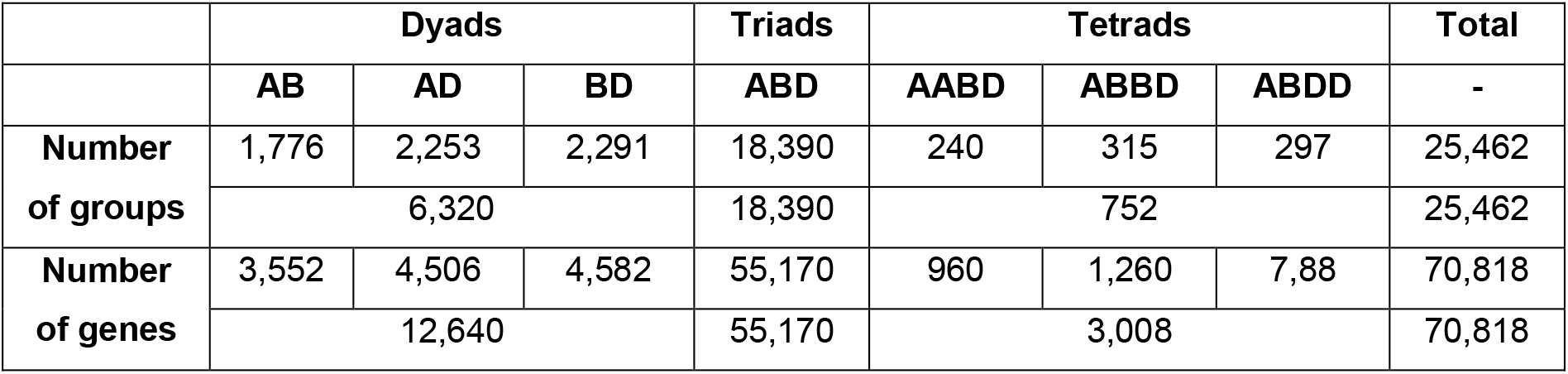
Number of groups and genes in the dyad, triad and tetrad categories

### Conservation of homoeoCNVs

To assess whether dyad genes were lost upon or after polyploidization or were already missing in the progenitor genomes, we searched for orthologs in diploid and tetraploid genomes: *Triticum urartu* (AA-genome), *Triticum dicoccoides* (AABB-genome) and *Aegilops tauschii* (DD-genome). For the A-missing copies (*i.e*. BD dyads), we estimated that approx. 40.7% and 19.0% were still present in diploid and tetraploid ancestor genomes, respectively (Data S1). For the B-missing (*i.e*. AD dyads) and D-missing ones (*i.e*. AB dyads), the estimates were of 20.6% and 27.7% still present in tetraploid and DD-diploid ancestor genomes, respectively. These results revealed that most of the genes of the dyad category were already absent from the diploid and tetraploid progenitors and that roughly 450 genes were lost on each subgenome at each step of polyploidization (498 A genes from *T. urartu* to *T. dicoccoides*; 434 A genes from *T. dicoccoides* to *T. aestivum*; 465 B genes from *T. dicoccoides* to *T. aestivum*; 493 D genes from *Ae. tauschii* to *T. aestivum*). Similarly, for tetrads, the majority of genes duplicated in the hexaploid wheat genome were also found in two copies in the diploid or tetraploid genomes. Indeed, 65.8% and 76.3% of A-duplicates were found in two copies in *T. urartu* and *T. dicoccoides*, respectively, 67.3% of B-duplicates in *T. dicoccoides* and 83.2% of D-duplicates in *Ae. tauschii*. Overall, we estimated that 172 genes were duplicated upon hexaploidization, 29.1% on the A-genome, 52.9% on the B-genome and 18.0% on the D-genome. However, one cannot exclude that the absence of a gene is due to an intraspecific polymorphism.

To investigate the conservation of dyad, triad and tetrad genes in other bread wheat accessions, we mined for presence-absence variations (PAVs) of genes in the genome of 16 resequenced wheat accessions (Montenegro et al., 2017). Out of the 70,818 genes, we identified 2,270 putative PAVs representing 3.2% of the dataset (Data S1). Consistent with the percentage of genes duplicated upon hexaploidization, the B-subgenome appeared to be more subject to variations (47.0% of all PAVs) than the A- and D-subgenomes (30.9% and 22.1%, respectively). When analysing each category of homoeogroups individually, only 1.8% of triad genes were affected by PAVs whereas they accounted for 7.9% and 9.7% of dyads and tetrads, respectively. Interestingly, for tetrads, 74.1% of PAVs appeared to affect one of the duplicated homoeologs.

Finally, to look at the conservation of dyad, triad and tetrad genes in other plants, we used the orthologous relationships with *Arabidopsis thaliana*, *Sorghum bicolor*, *Zea mays*, *Oryza sativa*, *Brachypodium distachyon* and *Hordeum vulgare* determined by the International Wheat Genome Sequencing Consortium (2018). The overall percentage of orthologs found for our 25,462 groups ranged from 52.8% in *A. thaliana* to 75.0% in *B. distachyon* (Table S2). These proportions were consistent with the phylogenetic distance, the most distant species sharing the lowest number of orthologs, with the notable exception of barley, consistent with the a lower BUSCO score indicating the completeness of genome assembly, gene set and transcriptome calculated by the IWGSC (2018). When analysing each category of homoeogroups separately, triad genes were found to be the most conserved (Figure 1). Indeed, the proportion of orthologous genes ranged from 58.4% in *A. thaliana* to 82.1% in *B. distachyon*. In tetrads, this proportion ranged from 37.6 to 56.4%. The least conserved genes were dyad ones with 32.0% of orthologs in *A. thaliana* and 48.3% in *B. distachyon*.

**Figure 1.**
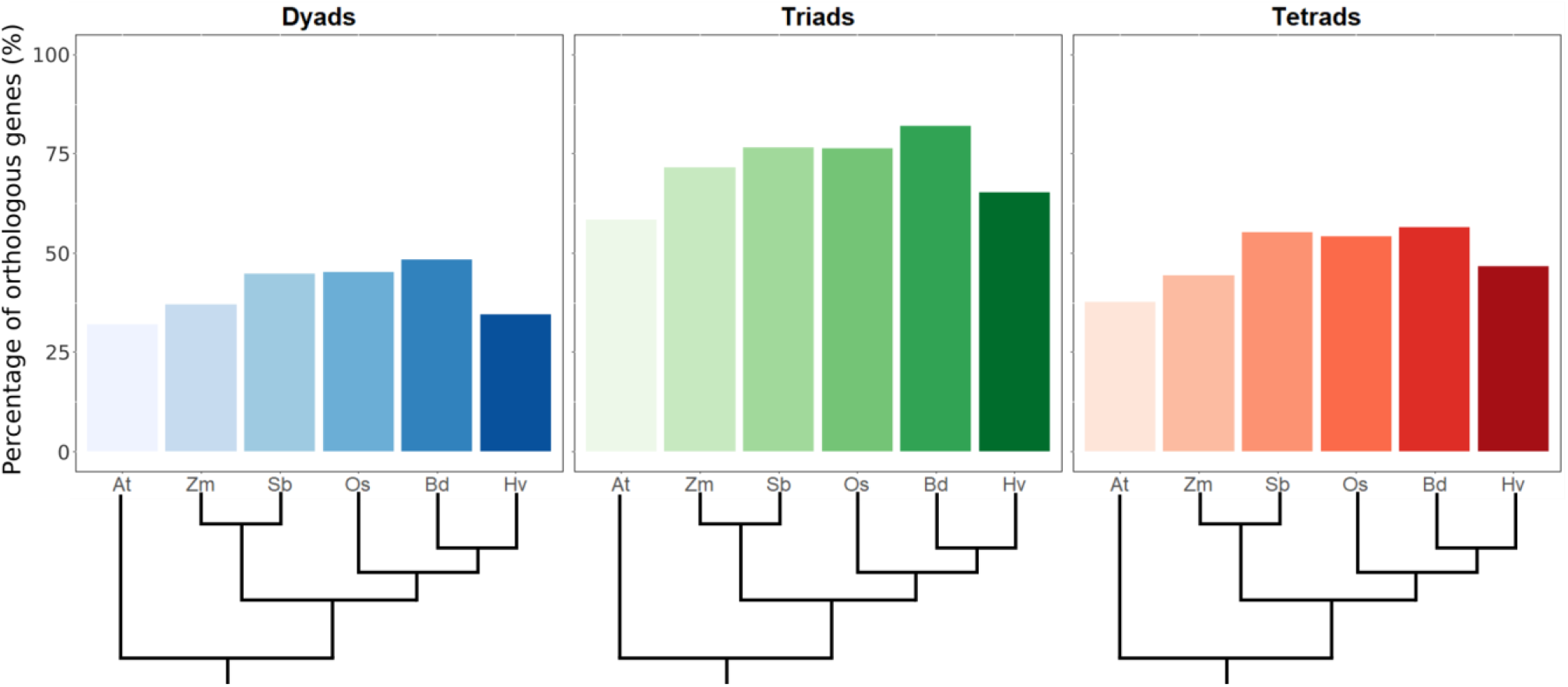
Conservation of genes in different plant genomes according to their category. For each category, the percentage of orthologs found in the *A. thaliana* (At), *Z. mays* (Zm), *S. bicolor* (Sb), *O. sativa* (Os), *B. distachyon* (Bd) and *H. vulgare* (Hv) genomes are given. Dyads are in blue, triads in green and tetrads in red.

### Distribution of homoeoCNVs along wheat chromosomes

Previous studies revealed a partitioning of the wheat genome based on different structural and functional features, including the recombination rate, gene and transposable element (TE) densities, gene expression breadth, histone modifications, as well as gene and TE structural variation rate (Choulet et al., 2014; De Oliveira et al., 2020; International Wheat Genome Sequencing Consortium, 2018; Pingault et al., 2015). Consequently, chromosomes can be divided into five chromosomal compartments: the short arm distal R1, the short arm proximal R2a, the centromeric-pericentromeric C, the long arm proximal R2b and the long arm distal R3 regions. To investigate the distribution of HomoeoCNVs in the light of chromosome partitioning, we analyzed the proportions of each category (dyads, triads, and tetrads) in the proximal (R2 and C) and distal (R1 and R3) regions of the chromosomes (Figure 2A and Table S2). We observed that triad homoeologs were more abundant in proximal than in distal regions: 64.3% *vs*. 35.7%, respectively. The opposite pattern was found for dyad genes with 62.2% located in distal regions and 37.8% in proximal regions. For tetrad genes, 57.6% were in distal and 42.4% in proximal regions.

**Figure 2.**
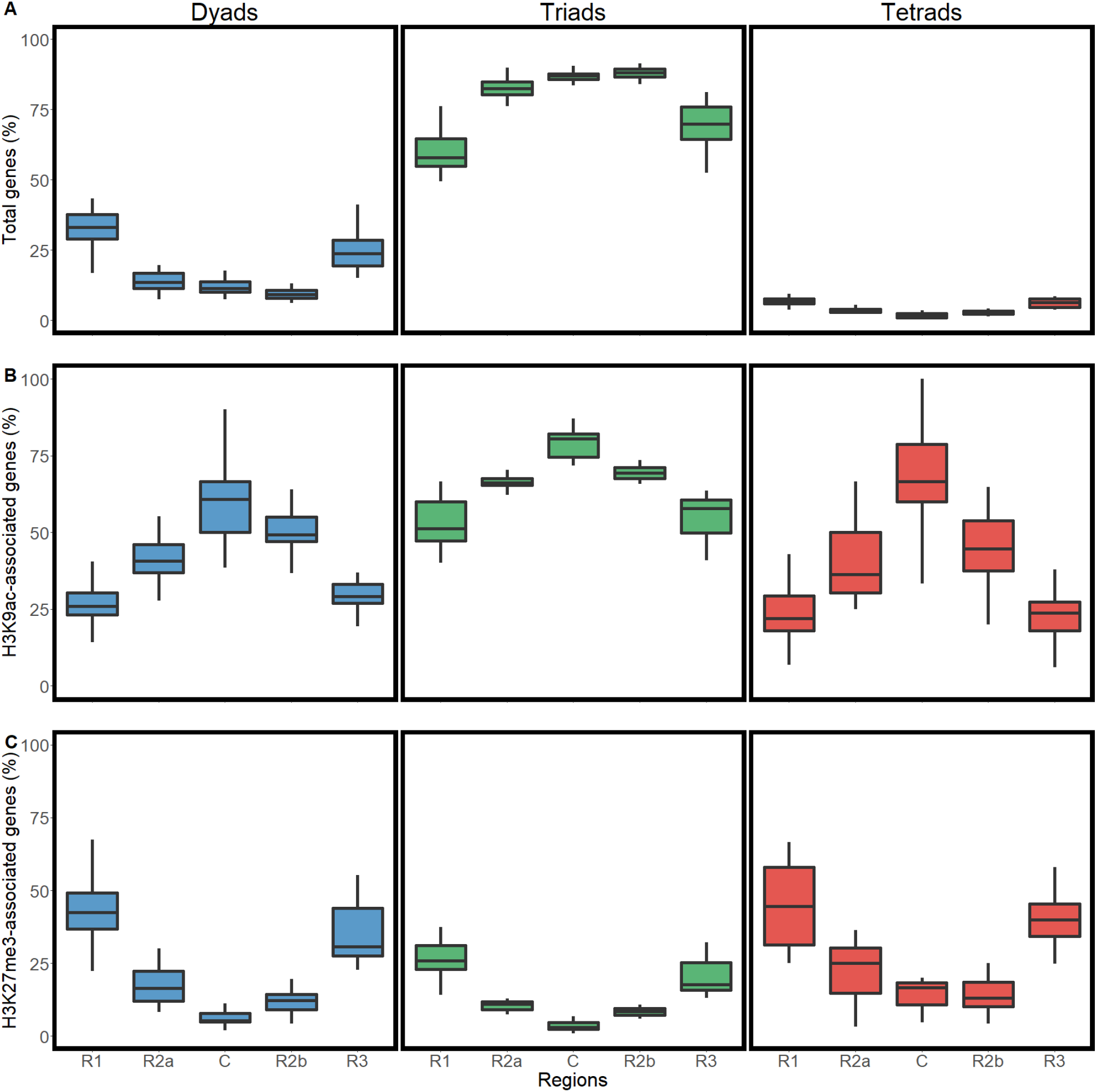
Distributions of genes of the different categories in the five regions of the wheat chromosomes. (A) percentage of genes of a given category in a given region according to the total number of genes in this region. (B) percentage of H3K9ac-associated genes according to the total number of genes from the same category in a given region (C) percentage of H3K27me3-associated genes according to the total number of genes from the same category in a given region. R1, R2a, C, R2b and R3 are the five chromosomal regions. Dyads are in blue, triads in green and tetrads in red.

At the chromosome scale, 95.6% of triads had their three genes located on homoeologous chromosomes (Figure 3). Out of the remaining 4.4%, 2.8% were found to have a mosaic distribution between chromosomes 4B, 4D and 5A or 4A, 5B and 5D, as a result of the structural evolution of the chromosomes 4A and 5A that have experienced inversions and translocations (Dvorak et al., 2018; Hernandez et al., 2012).

For dyads, 83.1% were found on homoeologous chromosomes and 3.3% showed a mosaic between chromosomes 4 and 5. For tetrads, the proportion of conserved homoeologous locations was much lower (73.1%) whereas that of mosaic distributions related to chromosome 4A evolutionary history was similar (3.2%).

As the boundaries of these regions are conserved between homoeologous chromosomes (International Wheat Genome Sequencing Consortium, 2018), we wondered to what extent the different gene copies of the same homoeologous group were located in the same regions (Figure 3). For triads, 90.7% of the genes belong to groups showing conserved locations for the three copies, with 30.8% being exclusively in distal regions and 59.9% being exclusively in proximal regions, confirming the high level of collinearity between A, B, and D. Only 9.4% of triad genes showed a variable location (mosaic distribution) between A, B, and D. For dyads, 57.0% of the homoeologs were exclusively located in distal regions, 32.7% were only in proximal and 10.3% were located in two different regions. For tetrads, 47.1% of the genes belong to groups having all their copies located exclusively in distal regions, 32.2% located exclusively in proximal regions and 20.7% with a mosaic distribution. The higher proportion of mosaic distribution for the tetrad category is explained by dispersed duplications that represented 36.6% of duplicated genes, of which 19.8% showed inter-chromosomal duplications.

**Figure 3.**
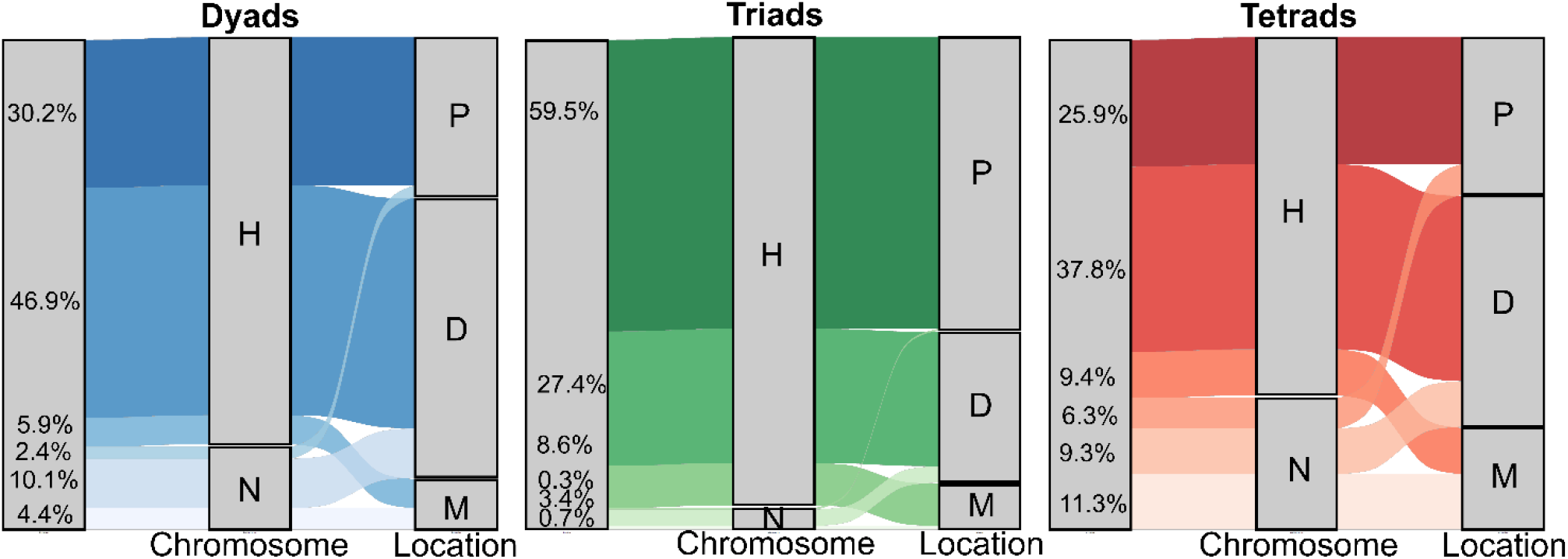
Alluvial plots of the different categories according to their location on chromosomes. For each category, the left-hand bar represents to whole set of genes, the central bar represents the position on homoeologous (H) or non-homoeologous (N) chromosomes; the right- and bar represents the location, either proximal for all copies of a group (P), distal (D) or a mosaic of proximal and distal genes (M). The number indicated in the left- and bar are the percentages of each class. Dyads are in blue, triads in green and tetrads in red.

### Functions of homoeoCNVs

Gene Ontology (GO) enrichment analysis revealed that dyads, triads and tetrads were involved in different biological processes. Indeed, triads were associated with basic cell processes such as transport, protein folding or DNA repair, replication and recombination. In contrast, tetrad and dyad genes were enriched in GO terms such as protein phosphorylation, oxidation-reduction processes, and response to fungus and oxidative stress (Table S3). Analysing the distal and proximal regions separately reached the same results, demonstrating that the GO enrichment was not only related to the preferential chromosomal location of the different categories (data not shown).

We expanded the analysis using the functional annotation of these genes to search for putative enrichment in protein functions (International Wheat Genome Sequencing Consortium, 2018) (Data S1). F-box family proteins appeared to be the most abundant family in dyads and tetrads, comprising 7.9% and 6.0% of genes, respectively, while it represented 2.1% of triads. Similarly, consistent with GO enrichment analyses, disease resistance associated genes such as NLR, RLK, BTB/POZ-domain or ankyrin represented 9.7% of dyads and 7.8% of tetrads but only 2.9% of triads. Among other functions enriched in dyads and/or tetrads compared to triads were oxidation-reduction processes-associated proteins such as peroxidases, Cytochrome P450 and glutathione S-transferases.

### Expression of homoeoCNVs

To evaluate expression differences between the three categories of homoeologs, we used a gene expression atlas covering the whole plant development in controlled non-stressed conditions (Pingault et al., 2015; Ramirez-Gonzalez et al., 2018). We found detectable expression (TPM values > 0.5) in at least one out of 15 tissues for 61,680 homoeologous genes from our dataset (87.1%) (Table S2 and Data S1). The proportion of expressed genes was slightly higher on the D-genome genes (87.7%) than on the A- and B-genomes (86.6% and 86.9%, respectively; *χ*^2^ p-value < 0.01). The percentage of expressed genes varied between categories too: 91.9% for triads (50,710 genes), 69.9% for dyads (8,838 genes) and 70.9% for tetrads (2,132 genes). In addition, we observed intra-category differences. For dyads, the AB-groups contained significantly fewer expressed genes (66.2%) than the BD- (70.5%) and AD-groups (72.2%) (χ2 p-value < 0.01). For tetrads, at a *χ*^2^ p-value of 1%, no significant differences were observed between groups. Interestingly, while in dyads and triads, the homoeologous genomes tended to have similar proportions of expressed copies, the duplicated-genome copies of tetrads displayed fewer expressed genes (χ^2^ p-value < 0.01; Table S2).

After discarding 7,179 outliers (649 dyad, 6,375 triad and 155 tetrad genes), we investigated the mean expression level and expression breadth (*i.e*. the number of tissues in which genes were expressed) of 54,501 expressed genes: 8,189 dyad, 44,335 triad and 1,977 tetrad genes. We found that triad genes were expressed at a higher level (mean = 5.9 TPM) and a higher breadth (10.4 tissues) than dyad (mean expression level = 5.2 TPM; mean expression breadth = 7.1) and tetrads (mean expression level = 5.2 TPM; mean expression breadth = 6.9) genes (Figure 4; Wilcoxon test p-value < 2.2e-16). To rule out the possibility that differences in expression level and breadth between dyads, triads and tetrads was only related to their chromosomal location, we divided the three categories in two sub-classes corresponding to their location, either proximal (R2 and C) or distal (R1 and R3). For all categories, genes located in proximal regions were expressed at significantly higher breadth than those in distal regions, confirming the impact of gene position on its expression. However, in general, triad genes were expressed at higher level and breadth than dyad and tetrad genes located in the same compartment (distal or proximal) (data not shown). This showed that the higher expression observed for triads may not only be due to their chromosomal location but to other factors. In tetrads, as for the proportion of expressed genes, the duplicated genome copies were less expressed than the non-duplicated ones, with lower expression breadth (6.6 *vs*. 7.1) and level (4.7 *vs*. 5.6; Wilcoxon test p-value < 0.01).

**Figure 4.**
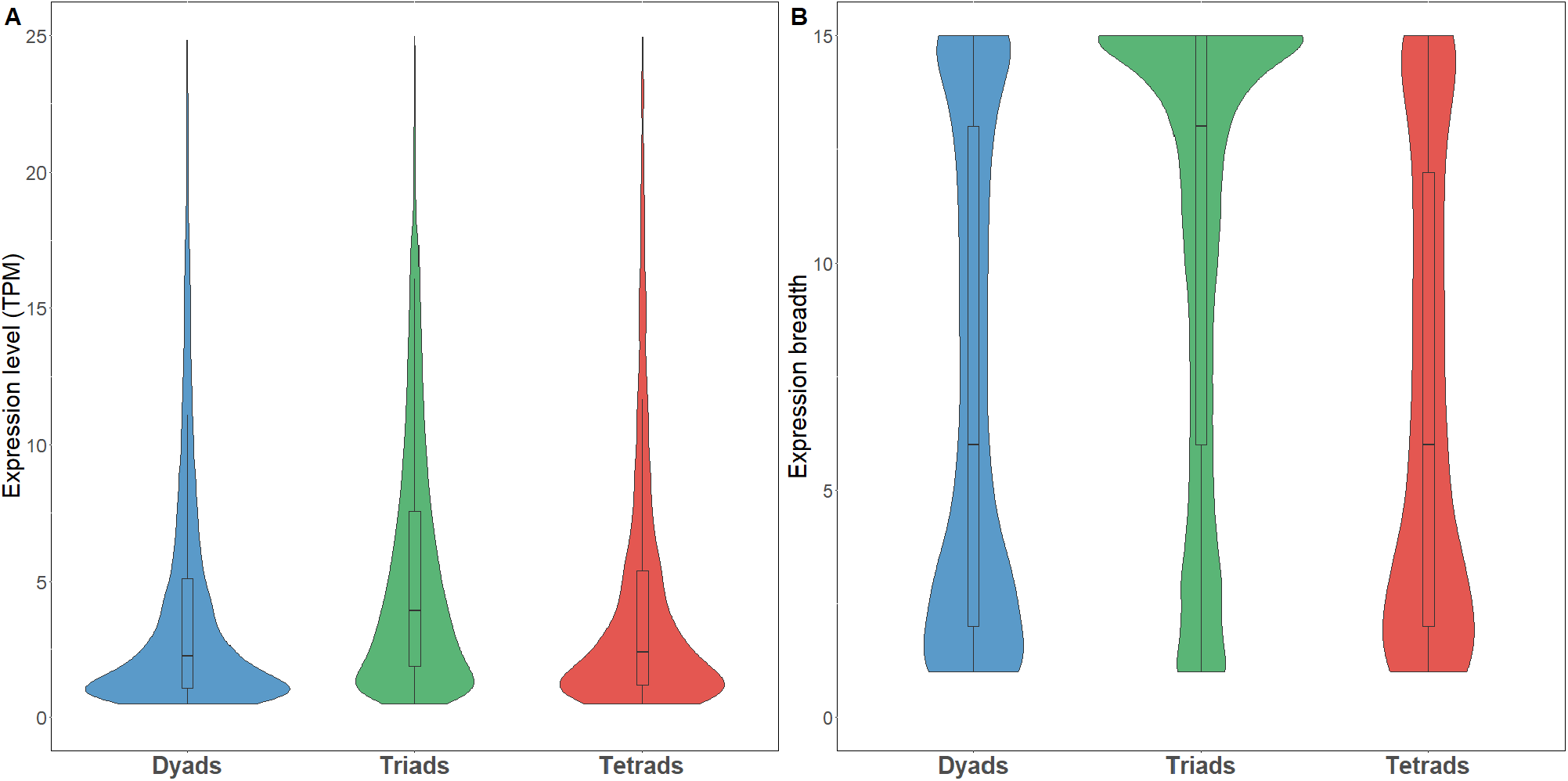
Expression level (A) and breadth (B) of the different categories. Expression level in TPM; expression breadth in number of conditions. Dyads in blue, triads, in green and tetrads in red.

### Relative contribution of each copy to the expression of the overall homoeologous group

To go further on expression analysis of our three categories of homoeologs, we calculated the relative contribution of each homoeolog to the overall group expression, for groups having at least one gene expressed. We then assigned each group to an expression bias category, as defined by Ramirez-Gonzalez et al. (2018): the balanced category with similar relative abundance of transcripts from each of the homoeologs, and the homoeolog-dominant or homoeolog-suppressed categories, classified based on the higher or lower abundance of transcripts from a given homoeolog with respect to those from the other(s) (Tables S2 and S3). For dyads, 64.0% of the groups were balanced, while 36.0% were dominant / suppressed. AB dyads appeared to be less frequently balanced than AD and BD dyads (60.1%, 66.6% and 64.2%, respectively; *χ^2^* p-value < 0.05). The expression breadth of balanced dyads was higher than that of suppressed / dominant ones (8.0 and 4.9, respectively) (Figure 5).

**Figure 5.**
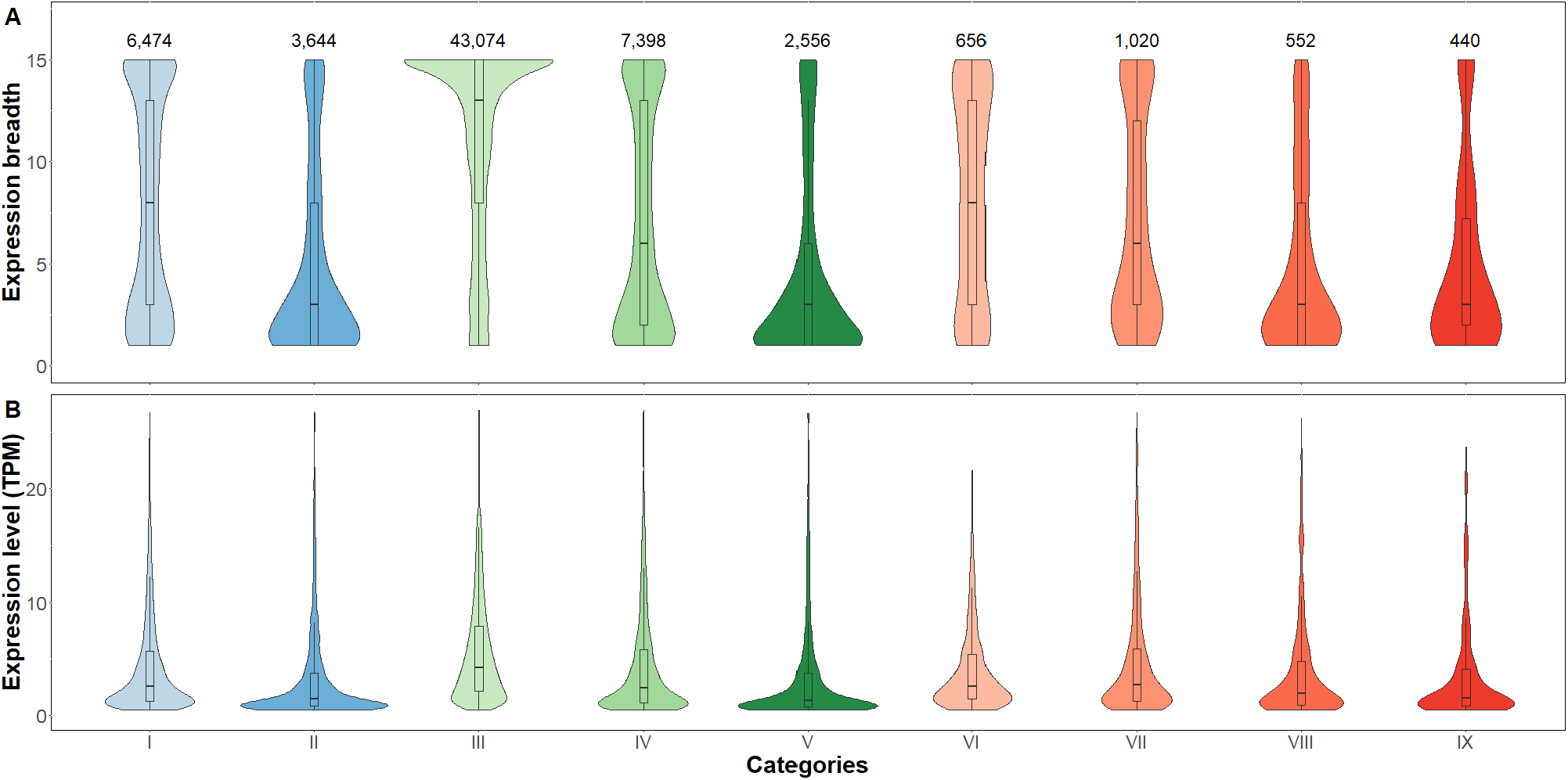
Expression breadth (A) and level (B) of the different categories according to their relative contribution classes. I: balanced dyads; II: Suppressed dyads; III: balanced triads, IV: Suppressed triads, V: Dominant triads, VI: balanced tetrads, VII: tetrads with one suppressed copy, VIII: tetrads with two suppressed copies, IX: tetrads with one dominant copy. The numbers above violin plots indicate the number of genes within the category.

For triads, an even higher proportion of balanced groups was observed (81.2%), while suppressed and dominant groups represented 14.0% and 4.8%, respectively. No difference was observed in the proportion of groups presenting a single-homoeolog dominance toward one sub-genome. Nevertheless, we observed a D-homoeolog suppression significantly less frequent (3.4%) than either A- or B-homoeolog suppression (5.3% and 5.2%, respectively; *χ^2^* p-value < 2.2e-16). As observed by Ramirez-Gonzalez and collaborators (2018), expression breadth decreased from balanced, to suppressed to dominant triads (11.1, 7.1 and 4.4, respectively) (Figure 5).

For tetrads, as expected from the greater number of gene copies, the pattern of relative contributions was much more complex. Balanced tetrads represented only 24.6% of the 667 groups having at least one expressed gene. The rest of the groups included 16.5% of groups with one copy dominant over the three others, 20.7% with two copies suppressed and 38.2% with one copy suppressed. It is worth noting that, for 74.5% of this latter, one of the two duplicates was suppressed. In addition, for ABDD tetrads, the duplication of a D-copy seems to have a similar impact on the suppression of A- and B-copies. By contrast, duplications of A-copies led to a slightly yet significantly higher proportion of B-copies than D-copies suppression (17.6% *vs*. 11.7%, respectively; *χ^2^* p-value < 0.01). A similar trend was observed in ABBD tetrads, where the B-copy duplication had greater impact on A-copy than D-copy suppression (15.7% *vs*. 11.2%, respectively; *χ^2^* p-value < 0.01). Interestingly, no significant difference was observed in terms of expression level between balanced tetrads and tetrads with one copy suppressed (5.2 TPM and 5.6 TPM, respectively) (Figure 5). The other tetrads displayed a significantly lower level (4.6 TPM for two suppressed copies and 4.6 TPM for one dominant copy; *χ*^2^ p-value < 0.01).

We then explored whether the different categories retain their homoeologous expression bias category across the five organs (root, leaf, stem, spike and grain) (Data S1). We found that 64.4% of balanced triads were also balanced (or not expressed) in the five organs, whereas, for dyads and tetrads, the proportions were 56.0% and 26.8%, respectively.

To complement this analysis, we investigated the divergence in spatial expression patterns. To this aim, we computed the binary expression (*i.e*. expressed or not) of each gene in the five different organs. This resulted in 32 binary expression clusters (0-0-0-0-0, 0-0-0-0-1, 0-0-0-1-1.). We then analysed each group to see whether genes from a given group belong to the same or divergent binary expression groups (Tables S2).

For triads, 65.3% had their three copies in the same cluster, among which 85.8% were expressed in all five organs. When analysing triads with one single divergent copy, we found a lower proportion of D-genome divergence, with 7.3% compared to 8.7% and 8.2% for the A and B-genomes, respectively (χ2 p-value < 0.01).

For dyads and tetrads, the proportion of groups having all the genes in the same binary expression cluster dropped to 45.7% and 21.1%, respectively. Interestingly, 21.9% of tetrads had one single divergent copy and in 71.2% of the cases, the divergent copy was one of the duplicates. The proportion of D-divergent ABDD tetrads was found to be lower (63.9%) even though the difference was not significant, probably due to the small sample size.

Finally, when considering only balanced groups, the percentage of groups having all their copies in the same binary expression cluster raised to 64.8% for dyads, 75.4% for triads and 46.3% for tetrads.

### Epigenetic status of homoeoCNVs

Epigenetic marks, and especially H3K9ac and H3K27me3 histone modifications, have been shown to be associated with differences in homoeolog expression patterns in triads (Ramirez-Gonzalez et al., 2018). These two marks have antagonist effects: H3K9ac is associated with open euchromatin and transcriptional activation whereas H3K27me3 is associated with facultative heterochromatin and transient transcriptional repression. To assess whether these marks may also be involved in the differences of expression patterns of dyads and tetrads, we analysed the presence of these two marks on the 70,818 genes from our dataset. We found 44,954 genes associated with H3K9ac and 15,357 with H3K27me3 (Data S1). After removing 3,809 genes that were associated with both marks, our dataset comprised 41,145 H3K9ac- and 11,548 H3K27me3-marked genes, *i.e*. 58.1% and 16.3% of all genes in the dataset, respectively (Table S2). These proportions differed according to the chromosomal locations: distal and proximal regions were enriched in H3K27me3 and H3K9ac genes, respectively, as shown previously (International Wheat Genome Sequencing Consortium, 2018) (Figures 2B and 2C).

As expected, H3K27me3 genes tended to be more often repressed than H3K9ac, with 31.5% and 3.5% of genes never expressed across the 15 tissues, respectively. We also found a higher expression breadth for H3K9ac genes (12.1 tissues) compared to H3K27me3 genes (2.8 tissues). When analysing gene expression in leaves at three-leaf stage (corresponding to ChIP-seq data), 85.1% of the H3K27me3 genes did not display any detectable expression while 83.2% of H3K9ac genes did.

When considering the three categories separately, we found differences in the proportion of the two marks. H3K27me3 was associated with 29.1% and 31.9% of dyad and tetrad genes, respectively, but only with 12.5% of triad ones (Figures 2B and 2C). By contrast, 64.7% of triad genes were marked by H3K9ac *vs*. 35.6% for dyads and 30.8% for tetrads.

Similar to what was observed previously, these proportions varied according to chromosomal regions (Figures 2B and 2C). However, triads were always more associated with H3K9ac and less with H3K27me3 than dyads and tetrads in the same chromosomal compartment. We also observed differences among tetrads. Indeed, the proportion of H3K9ac-associated genes increased from AABD to ABBD to ABDD (26.3%, 31.7% and 34.8%), while the opposite pattern was observed for H3K27me3 (34.7% for AABD, 31.7% for ABBD and 28.9% for ABDD). In addition, a significantly lower proportion of H3K9ac-associated genes was observed for the genome carrying the duplicated copies compared to the two others (27.7% vs. 33.9%; *χ*^2^ p-value < 0.01). For H3K27me3, the proportion was slightly higher yet not significantly (33.1 vs. 30.8%; (χ2 p-value > 0.01).

At the gene scale, balanced dyad, triad and tetrad genes had generally high H3K9ac and low H3K27me3 densities (Figure 6). For suppressed / non-suppressed and dominant / non-dominant groups, suppressed and non-dominant copies displayed higher H3K27me3 and lower H3K9ac than non-suppressed and dominant ones. Interestingly, the higher the number of suppressed copies in a group (from one to two in triads and from one to three in tetrads), the higher the H3K27me3 density, not only in the gene body but also into the upstream and downstream regions. This is consistent with the tight association of this mark with inactive promoters (Zhang et al., 2007).

**Figure 6.**
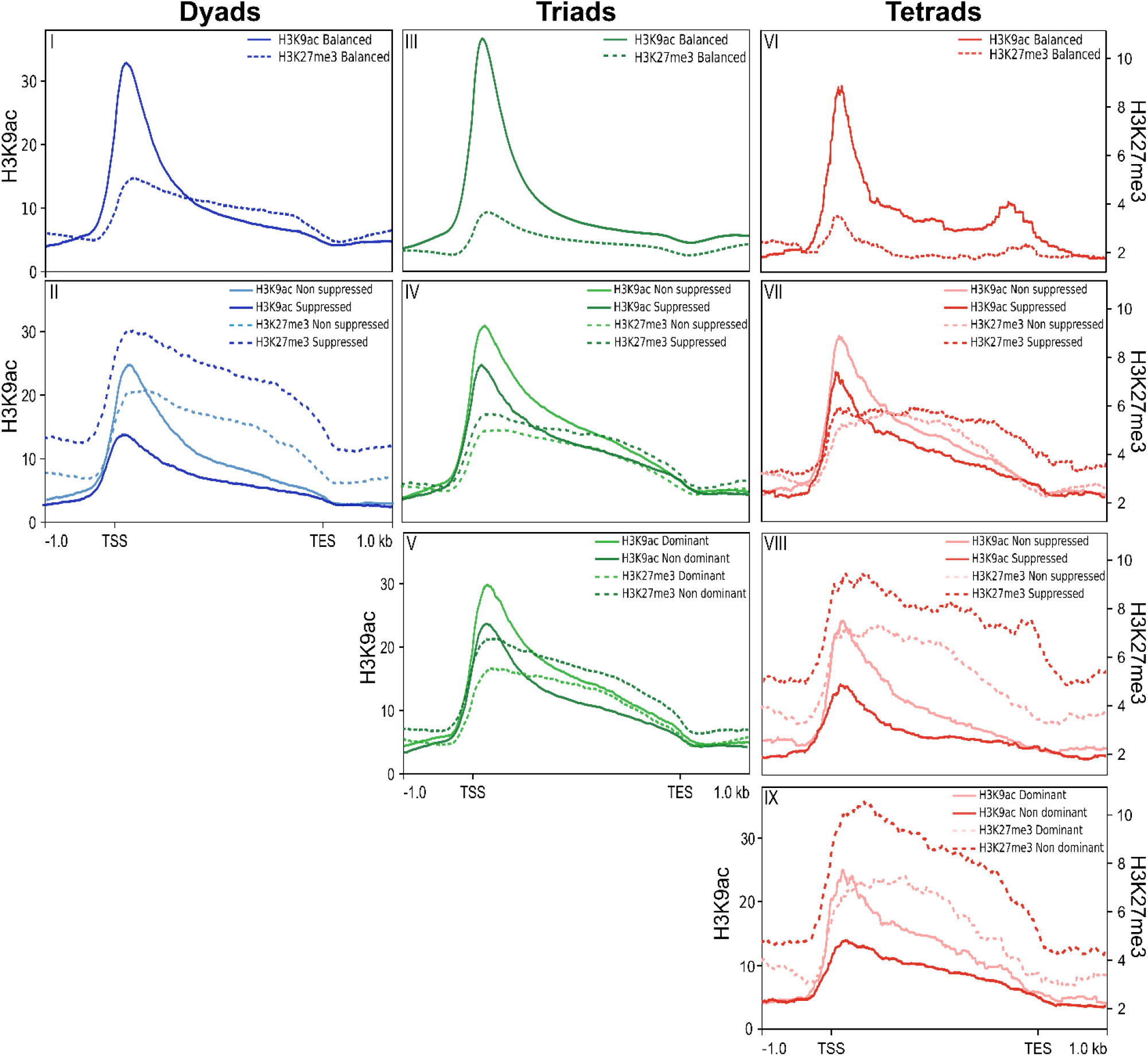
Epigenetic profiles of the different categories according to their relative contribution classes. Metagene profile for histone H3K9ac (solid lines) and H3K27me3 -dashed lines) marks from −1 kb upstream of the ATG to +1 kb downstream of the stop codon (normalized for gene length) for dyads (blue), triads (green) and tetrads (red) dominant / non-suppressed (light colours) and suppressed / nondominant (dark colours) copies. I: balanced dyads; II: Suppressed dyads; III: balanced triads, IV: Suppressed triads, V: Dominant triads, VI: balanced tetrads, VII: tetrads with one suppressed copy, VIII: tetrads with two suppressed copies, IX: tetrads with one dominant copy.

We then analysed whether genes from a group containing at least one gene associated with a mark tended to share the same epigenetic mark. For triads, 64.4% of the groups comprised three genes sharing the same mark, either H3K9ac (88.5%) or H3K27me3 (11.5 %). This percentage was lower (55.9%) for dyads (57.1% H3K9ac and 42.9% H3K27me3). For tetrads, only 28.1% of groups comprised four genes sharing the same mark. Nevertheless, this percentage raised to 52.8% when including groups with three copies sharing the same mark. Finally, we investigated the conservation of histone marks in two other species, *Zea mays* and *Oryza sativa*, for which H3K27me3 and H3K9ac data on a young leaf stage were available in the Plant Chromatin Database (Liu et al., 2018). Out of 12,995 and 12,006 groups containing at least one wheat H3K9ac-marked gene that had an ortholog in rice and maize, 10,541 (81.1%) and 7,604 (63.3%) were also marked by H3K9ac in these two species, respectively (Table S2). For H3K27me3, the conservation was lower with 47.8% and 56.4% of the groups containing orthologs also targeted by this mark in rice and maize, respectively.

Surprisingly, while strong differences were observed at the sequence orthology level between dyads, triads and tetrads, the conservation of histone marks was not so different between the three categories. For example, while triads were much more conserved with rice than dyads (76.4% *vs*. 45.1%, respectively), quite similar proportions of groups containing conserved histone-marked genes were observed (48.4% of triads and 46.7% of dyads for H3K27me3, 81.5% of triads and 79.1% of dyads for H3K9ac).

## Discussion

Wheat is an allohexaploid species originating from two successive and recent rounds of hybridization between three diploid species that were very similar in terms of chromosome number, genome size, TE content, gene content and synteny (International Wheat Genome Sequencing Consortium, 2018; Wicker et al., 2018). As a result, and considering that wheat is an autogamous homozygous species, it has long been considered that most of the genes were in three homoeologous copies. This perception started to change with the advent of the first draft assembly of the wheat genome sequence ((International Wheat Genome Sequencing Consortium, 2014). The reference sequence of the hexaploid wheat genome confirmed that a significant fraction of genes departed from this 1:1:1 ratio and that these so-called ‘triads’ represent less than one half of all wheat genes.

In a recent work, Ramirez-Gonzalez et al. (2018) characterized the transcriptome atlas of wheat with a focus on these triads. This analysis provided new insights into the relative contribution of homoeologous copies to the overall group expression and the possible role of epigenetic marks in establishing this pattern.

In this study, we extended this analysis to genes departing from the 1:1:1 ratio, and more particularly the homoeologous groups having undergone a single gene loss or duplication event. These so-called dyads and tetrads, collectively referred to as HomoeoCNVs, represented 17.8% and 4.2% of our HC gene datasets whereas triads represented 77.9%. These proportions differed from those reported by the IWGSC (2018) as we focused our analysis on HC genes while the filtered dataset used by the IWGSC consisted of both HC and LC genes and took into account all categories, including genes in N:N:N ratio and not only dyads, triads and tetrads.

Because they have been kept in a strict 1:1:1 ratio through the course of evolution, triads are likely to correspond to highly conserved and evolutionary constrained genes. By contrast, dyads and tetrads have been either deleted or duplicated in the hexaploid wheat genome or in its diploid and tetraploid progenitors. We therefore suggest that triads mainly correspond to housekeeping genes and are part of the core genome, while dyads and tetrads belong to the dispensable genome of wheat. Several findings support this hypothesis.

First, we found that triads were more conserved in other plant genomes than HomoeoCNVs. By contrast, dyads and tetrads were found to be less conserved not only in distant plant genomes such as *A. thaliana*, *Z. mays*, *O. sativa*, *S. bicolor* or *B. distachyon* but also in the *Triticum* / *Aegilops* species, as most of these genes were already missing or duplicated in the wheat progenitors. In addition, HomoeoCNVs were also enriched in PAVs in a panel of 16 hexaploid wheat accessions. Previous studies in soybean, rice and *B. distachyon* demonstrated that core genes tend to have a higher percentage of homologs in other species than dispensable ones (Gordon et al., 2017; Li et al., 2014; Zhao et al., 2018). In wheat, this difference in terms of gene conservation is consistent with the genomic distribution of the different categories and the chromosome partitioning (Choulet et al., 2014; Daron et al., 2014; Darrier et al., 2017; Glover et al., 2015; International Wheat Genome Sequencing Consortium, 2018). Indeed, we showed that triads were more abundant in the low-recombination proximal regions. By contrast, dyads and tetrads were enriched in distal regions where differential TE content and recombination rate have likely driven gene duplications and deletions (Akhunov et al., 2003; Dvorak and Akhunov, 2005; Feldman et al., 2012; Reams and Roth, 2015; Zhang, 2003).

We also found that triads were expressed at higher level and breadth, while dyads and tetrads tend to be more specific to some tissues or developmental stages. In *B. distachyon*, core genes tend to be expressed at a higher level and more broadly than dispensable genes (Gordon et al., 2017). Choulet et al. (2014) and Pingault et al. (2015) reported on the physical partitioning of wheat genes, with highly and constitutively expressed genes being mainly located in proximal regions and genes expressed at lower level and breadth in distal ones. However, by analysing distal and proximal regions separately, we showed that triads were always more expressed than dyads and tetrads whatever their position on the chromosome, which ruled out the possibility that the differences in expression patterns were only related to the chromosomal positions. Conversely, this difference can at least partly be explained by the epigenetic pattern of the categories of homoeologous genes. Indeed, triads were enriched in H3K9ac active euchromatin mark whereas dyads and tetrads were enriched in H3K27me3, a repressive mark related to facultative heterochromatin (Wiles and Selker, 2017). This differential association with active or repressive histones marks have already been reported in other species, such as potato where CNV frequency increased in genes lacking histone marks associated with permissive transcription (Hardigan et al., 2016). In wheat, we showed recently that genes affected by intra- and interspecific copy number variations were enriched in H3K27m3 (De Oliveira et al., 2020).

Finally, dyads and tetrads were enriched in functions associated with environmental and defence responses, a common feature of most plant dispensable genomes (Golicz et al., 2016; Gordon et al., 2017; Hurgobin et al., 2018; Li et al., 2014; McHale et al., 2012; Schatz et al., 2014). In particular, we found a higher proportion of genes associated to oxidation-reduction process that are known to be related to reactive oxygen species and putatively to biotic (pathogens) and abiotic (heavy metals, salt...) stress response mechanisms (Gullner et al., 2018; Mir et al., 2015; Mittler et al., 2004; Veith and Moorthy, 2018). Disease resistance-associated families, such as NLRs, RLKs, ankyrin repeat or BTB/POZZ domain-containing proteins were also found in higher proportions in dyads and tetrads than in triads (Sun et al., 2020; Wang et al., 2020; Ye et al., 2017; Zhang et al., 2019).

We then examined intra-categories differences to investigate the possible impact of polyploidization on both core and dispensable genomes in wheat. Indeed, polyploidization is usually followed by a post-polyploid diploidization (PPD) process that tends to revert the polyploid genome into a quasi-diploid one (Mandáková and Lysak, 2018). PPD is accompanied by several mechanisms including gene neo/subfunctionalization, activation of transposable elements, epigenetic reprogramming and genome fractionation. Genome fractionation is a long-term process involving the loss of redundant genes and/or noncoding regulatory elements (Cheng et al., 2018). While it has been observed in several species and seems to be a common mechanism, differences have been observed according to the type of whole genome duplication (Garsmeur et al., 2013). In allopolyploids or paleo-allopolyploids such as *Arabidopsis thaliana*, maize (*Zea mays*), Chinese cabbage (*Brassica rapa*) and *Brassica oleracea*, duplicated genes are lost preferentially from one parental genome (biased fractionation) and the subgenome having retained the highest number of genes is more expressed (genome dominance) (Liu et al., 2014; Schnable et al., 2011; Wang et al., 2011). By contrast, in autopolyploids or paleo-autopolyploids, such as poplar (*Populus trichocarpa*) and pear (*Pyrus bretschneideri*), subgenome dominance is absent and genes tend to be evenly lost between the two subgenomes (Li et al., 2019; Liu et al., 2017). In wheat, some rapid changes following polyploidization have been reported, including chromosomal rearrangements, epigenetic changes or TE-related shift in centromere position (Badaeva et al., 2015; Dvorak et al., 2018; Jiao et al., 2018; Li et al., 2013; Liu et al., 2009; Shaked et al., 2001; Zhao et al., 2019). However, whether the wheat genome experiences subgenome dominance or biased fractionation is still a matter of debate. Different analyses reached contradictory results (El Baidouri et al., 2017; International Wheat Genome Sequencing Consortium, 2018; Pont and Salse, 2017).

Consistent with what was observed on a filtered set of 181,036 genes comprising both HC and LC genes (International Wheat Genome Sequencing Consortium, 2018), the number of genes analysed in our study was highly similar between subgenomes, with 23,411 on A, 23,524 on B and 23,883 on D. These similar proportions can be partly explained by the fact that the vast majority of these groups (75.9%) corresponded to triads, with one copy on each of the subgenomes. Such a high percentage of genes that are still present on the A, B and D-genomes demonstrate that no massive gene loss occurred upon polyploidization. Homoeologous groups that have lost one copy, *i.e*. dyads, represented 17.8% of our dataset. This category might reflect post-polyploidization gene loss. Interestingly, a lower number of AB-dyads (1,776) was observed compared to AD- or BD-ones (2,253 and 2,291, respectively). However, the analysis of the diploid and tetraploid ancestors suggested that only approx. 450 genes were lost on each subgenome at each step of polyploidization. This similar number of lost genes reveals the absence of a biased fractionation in wheat. Nevertheless, while no bias was found in terms of gene loss, we noticed subtle differences between genomes at the transcription and epigenetic levels.

The majority of triads displayed a balanced contribution of each copy to the overall group expression (81.2%). They also showed a high proportion of homoeologous genes having the same binary spatial expression (65.3%) and sharing the same histone mark (69.3%). However, we observed a slightly higher proportion of D-genome expressed genes compared to A and B, together with a lower proportion of D-suppressed triads, and a lower proportion of D divergent copies. This suggests a lower repression or subfunctionalization of D-genome homoeologs compared to their A and B counterparts. While these are very faint differences, they might explain the subtly yet significantly higher relative abundance of the D-subgenome transcripts (33.7%) compared to the A (33.1%) and B (33.3%) reported by Ramirez-Gonzalez and collaborators (2018).

Similarly, little differences were observed in dyads. They were mainly balanced (64.0%), with most of the homoeologs associated with the same epigenetic mark (65.1%). In addition, we found no bias in terms of dominance of one genome over the other. However, AB-dyads tended to be slightly less numerous and less expressed than the AD- and BD-ones, these two latter being similar on several aspects.

The pattern was much more complex for tetrads, with fewer balanced groups (24.6%) that might be explained by a higher proportion of mosaic patterns of epigenetic marks (71.9%). The overall higher proportion of H3K27me3- and lower proportion of H3K9ac-associated genes, especially on the genome carrying the duplicated copy, were likely related to subfunctionalization of retained paralogs as suggested by Makarevitch et al. (2013) and Berke et al. (2012). According to their mean expression level and breadth, as well as the epigenetic pattern, ABDD tetrads were more comparable to AB-dyads than to other tetrads. Indeed, the A and B copies of both ABDD tetrads and AB dyads were more associated with H3K9ac and less with H3K27me3, while the opposite pattern was found for those of AABD and ABBD tetrads. In addition, an extra copy on the B-genome (ABBD tetrads) appeared to have a much stronger impact on the A-genome than on the D-genome, while an extra D-copy had a similar impact on A and B in ABDD tetrads.

Based on these different results, we propose that the D-subgenome homoeologous genes are less repressed than the two others, and conversely, that their presence, either as single or duplicated genes, had a limited impact on the A- and B-copies. Differences observed between subgenomes are likely related to the D-genome more recent hybridization with the AABB tetraploid genome progenitor. This resulted in two successive PPD waves, that had impacted A and B more significantly since they spent more time together. However, unlike most of the allopolyploid species, subgenome dominance and biased fractionation are absent in hexaploid wheat. Indeed, while originating from the hybridization of three distinct species, the diploid donor genomes were very similar in terms of gene and TE contents prior to polyploidization. Consequently, individual genes, rather than subgenomes, experienced stochastic differences over longer periods of time, resulting in retention of the majority of WGD duplicates. In this regard, while being an allohexaploid species, wheat somehow resembles more to an autopolyploid in terms of evolutionary fate, as already observed in other paleo-allopolyploids such as soybean (*Glycine max*) and cucurbits (*Cucurbita maxima* and *Cucurbita moschata*) (Sun et al., 2017; Zhao et al., 2017).

## Supplemental material

**Table S1. Definition of homoeolog expression bias categories**

**Table S2. Summary of dyads, triads and tetrads features**

This table contains analyzed data including the number of homoeologous groups and genes, Number and percentage of genes in distal and proximal location, Number and percentage of expressed genes in the three subgenomes and in different conditions, Number and percentage of expressed genes without outliers, Expression level without outliers, Expression breadth without ouliers, Number of orthologous genes found in, Percentage of orthologous genes found in other species, Number and percentage of H3K27me3-associated genes, Number and percentage of H3K9ac-associated genes, Percentage of orthologs sharing the same histone mark, Relative contribution categories of dyad genes, Relative contribution categories of triad genes, Relative contribution categories of tetrad genes, Organ binary expression numbers, Organ binary expression percentage

**Table S3. Gene Ontology enrichment results**

For dyads, triads and tetrads, the most significant GO terms (according to the topGoFisher test, p-value < 0.01) are listed, with their relative statistics.

**Data S1. Homoeologous gene data**

This table contains all homoeologous gene data including Gene ID; Genome on which gene is located; Chromosome on which gene is located; Start position of gene on IWGSC RefSeqv1.0; Stop position of gene on IWGSC RefSeqv1.0; Assignment to one of the five chromosomal regions (R1, R2a, C, R2b a,d R3); Assignment to a proximal (P) or distal (C) compartment; Homoeologous group category (Dyad, triad or tetrad); Detailed homoeologous group category; Homoeologous group ID; Chromosomes where the homoeologous copies are located; Expression breadth (in number of conditions, from 0 to 15); mean expression level across the 15 tissues (in TPM); mean expression level in root (in TPM); mean expression level in leaf (in TPM); mean expression level in stem (in TPM); mean expression level in spike (in TPM); mean expression level in grain (in TPM); mean expression level in leaf at three-leaves stage (in TPM); Identified as outlier (TRUE) or not (FALSE); Relative contribution category in details; Relative contribution category summary; Relative contribution category in brief; Relative contribution category in details in root; Relative contribution category in details in leaf; Relative contribution category in details in stem; Relative contribution category in details in spike; Relative contribution category in details in grain; Binary expression cluster; Presence of the H3K27me3 mark; Presence of the H3K9ac mark; Epigenetic profile; Identified as PAV (ys) or not (no) in 16 wheat accessions; A-missing copy in hexaploid wheat found in Triticum urartu genome; A-missing copy in hexaploid wheat found in Triticum dicoccoides A-genome; B-missing copy in hexaploid wheat found in Triticum dicoccoides B-genome; D-missing copy in hexaploid wheat found in Aegilops tauschii genome; Oryza sativa orthologous gene ID; Zea mays orthologous gene ID; Brachypodium distachyon orthologous gene ID; Arabidopsis thaliana orthologous gene ID; Hordeum vulgare orthologous gene ID; Sorghum bicolor orthologous gene ID; H3K9ac-associated Oryza sativa orthologous gene ID; H3K27me3-associated Oryza sativa orthologous gene ID; H3K9ac-associated Zea mays orthologous gene ID; H3K27me3-associated Zea mays orthologous gene ID; Putative annotated function

## Acknowledgments

We acknowledge Hélène Rimbert and Philippa Borrill for their assistance. C.J. was supported by Région Auvergne and the European Regional Development Fund (SRESRI 2016).

## Abbreviations

GO: Gene ontology
HC: High Confidence
HomoeoCNV: Homoeologous Copy Number Variation
LC: Low Confidence
MYA: Million Years Ago
PAV: Presence Absence Variation
PPD: Post-Polyploid Diploidization
TE: Transposable Element
TPM: Transcripts Per Million
WGD: Whole Genome Duplication

